# Predation Has Small, Short-Term, and in Certain Conditions Random Effect on the Evolution of Aging

**DOI:** 10.1101/2020.05.13.092437

**Authors:** Peter Lenart, Julie Bienertová-Vašků, Luděk Berec

**Affiliations:** Research Centre for Toxic Compounds in the Environment, Faculty of Science, Masaryk University, Kamenice 5, Building A29, 62500, Brno, Czech Republic; Department of Experimental Biology, Faculty of Science, Masaryk University, Kamenice 5, 62500 Brno, Czech Republic; Centre for Mathematical Biology, Institute of Mathematics, Faculty of Science, University of South Bohemia, Branišovská 1760, 37005 České Budějovice, Czech Republic; Czech Academy of Sciences, Biology Centre, Institute of Entomology, Department of Ecology, Branišovská 31, 37005 České Budějovice, Czech Republic

## Abstract

The pace of aging varies considerably in nature. Historically, scientists focused mostly on why and how has aging evolved, while only a few studies explored mechanisms driving evolution of specific rates of aging. Here we develop an agent-based model simulating evolution of aging in prey subject to predation. Our results suggest that predation affects the pace of aging in prey only if young, vivid animals are not much more likely to escape predators than the old ones. However, even this effect slowly vanishes when the predator diet composition evolves, too. Furthermore, evolution of a specific aging rate, in our model, is driven mainly by a single parameter, the strength of a trade-off between aging and fecundity. Indeed, in absence of this trade-off the evolutionary impacts of predation on the prey aging rate appear random. Our model produces several testable predictions which may be useful for other areas of aging research.

## Introduction

Aging, defined as an age-dependent increase in the mortality rate^1^, is a widespread biological phenomenon. However, it seems that some species do not age at all^2–5^ or at least do not age at the scale comparable with others. In addition, in species which do age, the actual pace of aging is quite diverse^2,6^, which suggests that the aging rate may be a trait malleable by evolutionary forces. A better understanding of the mechanisms driving the evolution of the pace of aging would certainly provide an important insight into how aging operates and how it can be modulated.

Classical evolutionary theories of aging postulate that aging is an inevitable by-product of evolution and serves no adaptive purpose^7–10^. There are two major problems with such views. First, the idea of inevitability of aging is directly contradicted by the existence of species with negligible senescence^3–5,11^. Second, if aging serves no adaptive purpose and is only detrimental, then non-aging (negligible senescence) is beneficial, suggesting that there should be a strong selective pressure driving evolution of non-aging in most long-lived species. However, non-aging has developed in only a minority of known species.

To cope with these problems, some authors proposed the existence of the so-called “aging program”^12–17^, together with mechanisms of its evolution^13,15,18^. Nevertheless, these attempts to explain evolution of aging are still heavily disputed^7,19^. Moreover, evolutionary models showing that aging can be adaptive explore mostly whether aging individuals can outcompete non-aging ones^20–23^, instead of an examination of factors driving the evolution of specific aging rates.

Acknowledging that aging is an inevitable result of damage accumulation, we have previously proposed that the actual pace of aging is adaptive and thus that in different conditions, different paces of aging may evolve, with negligible senescence being only an extreme case of very slow aging^24^. In addition, we have developed a mathematical model showing that under some conditions, sexual reproduction promotes aging over non-aging^25^. These results explain why negligible senescence is not common in nature but tell us nothing about inevitability of aging itself or mechanisms driving the evolution of specific aging rates. In this article, we develop a novel mathematical model that simulates evolution of the aging rate under various ecological scenarios. In particular, we ask how the presence of predators affects the aging rate in their prey, and how this may in turn shape predators’ consumption preferences for prey of different age. Also, we ask how these relationships are modulated by commonly assumed life history trade-offs, an aging rate-reproduction trade-off in prey and a searching effort-maintenance trade-off in predators.

## Methods

To address our questions, we develop an agent-based simulation model of a predator-prey interaction that allows all prey and predator individuals and hence their phenotypes to be modelled explicitly and their competitive, predatory and phenotypic dynamics to be followed over time. Time is discrete, with the time step corresponding to the age increment of 1 and with all relevant rates and probabilities defined on the per time step basis. We consider populations composed of *N* prey and *P* predators, both formed by (haploid) simultaneous hermaphrodites. All model parameters and variables are summarized in Table 1.

**Table 1:** Default parameter settings used in our simulation model.

| Parameter | Meaning | Value(s) |
| --- | --- | --- |
| $N$ | Fixed prey population size | 1000 |
| $P$ | Fixed predator population size | varies |
| $d_0$ | Per time step probability of dying at age 1 for prey individuals | 0.01 |
| $k_d$ | Rate at which mortality of prey individuals increases with age | trait |
| $b_0$ | Fecundity of prey individuals at age 1 | 1, 5 |
| $p_m$ | Probability of mutation upon offspring production | 0.5 |
| $\sigma_m$ | Variance in trait upon mutation | 0.01 |
| $w_{d0}$ | Minimum predator attack rate | 0.001 |
| $w_1$ | Maximum predator attack rate | 0.01 |
| $k_w$ | Flexibility in the predator attack rate | trait |
| $m$ | Per time step probability of dying for prey individuals | 0.02 |
| $e_0$ | Assimilated energy from prey of age 1 | 1 |
| $x$ | Trade-off strength in prey trait $k_d$ | 0, 0.01, 0.05 |
| $c$ | Trade-off strength in prey trait $k_w$ | 2 |
| $e_1$ | Energy needed to produce one predator offspring | 1 |
| $k_e$ | Marginal rate in assimilated energy with prey age | 0.1 |
| $k$ | Maintenance parameter in predators | varies |

### Prey demography

Prey individuals are characterized by age *a*, and their phenotypes are assumed to differ according to the mortality rate profile. We define the probability that a prey individual of age *a* dies during a single time step as

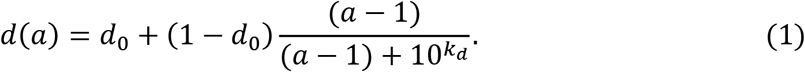

This function increases in a decelerating way from 0 < *d*_0_ < 1 for *a* = 1 to 1 as *a* grows large (Fig. 1a). Different prey phenotypes are distinguished by different values of the parameter *k_d_* which determines the rate at which mortality increases with age. Specifically, higher *k_d_* means less steep slope at *a* = 1 and slower approach of the limiting value 1, and hence slower aging (Fig. 1a). Evolution of the aging rate is thus in this setting equivalent to evolution of the parameter *k_d_* within the prey population. The reason for not introducing *k_d_* directly as the half-saturation constant in the above fraction but rather as an exponent of 10 is to more easily explore many orders of magnitude the half-saturation constant might admit and not to care for the lower bound of zero; it is just a technical assumption that however allows simulations to run more efficiently.

**Figure 1:**
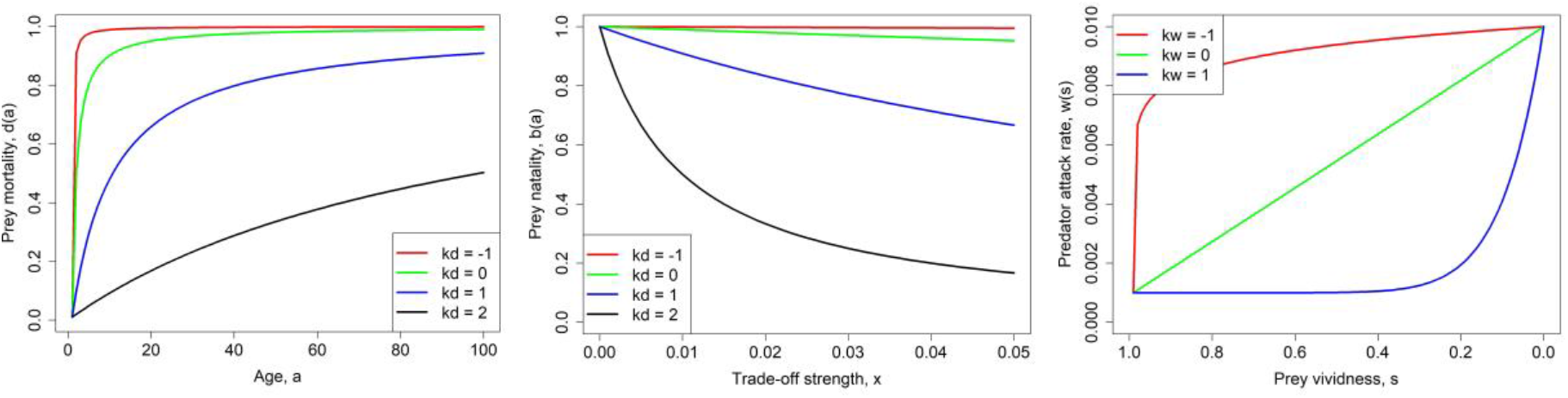
Functional relationships describing various model elements. (a, left) Prey mortality described by equation (1). (b, middle) Prey fecundity described by equation (2). (c, right) Predator attack rate of prey described by equation (3). Parameters: *b*_0_ = 1, *d*_0_ = 0.01, *w*_*d*0_ = 0.001, *w*_1_ = 0.01. We note that prey vividness decreases with age as *s*(*a*) = 1 – *d*(*a*).

Since slower aging likely does not come for free yet rather negatively impacts another individual trait, we assume that an increase in the aging rate parameter *k_d_* (i.e. slower aging) implies lower individual fecundity and vice versa, as suggested by the disposable soma theory^9,10^. There are many alternative ways of how to model this relationship. Denoting by *x* the strength of trade-off between the aging rate and fecundity, we use the following relationship (Fig. 1b):

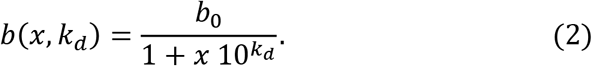

Offspring mortality in their first year of life is included in the parameter *b*_0_ and equations (1) and (2) are thus applicable to prey individuals of age *a* ≥ 1.

### Predation and predator demography

Predator population is assumed unstructured with respect to age but the attack rate of predators towards prey depends on the age of prey *a*. Specifically, ‘older’ prey individuals are assumed to have reduced abilities to escape predators. But ‘older’ has a relative meaning, as various prey individuals may age at different rates, so we should rather say that less vivid individuals are assumed to have reduced abilities to escape predators. Such a distinction between biological age and physical performance is also well supported by existing data: a recent study performed on a sample of 126 356 subjects shows that age estimated based on exercise stress testing performance is a better predictor of mortality than chronological age^26^. We thus consider the prey survival probability *s*(*a*) = 1 – *d*(*a*) as a proxy of prey vividness. Note that different prey phenotypes may have the same vividness at different ages which is exactly what we aim here for. We model the predator attack rate of prey as a decreasing function of the prey vividness *s*(*a*):

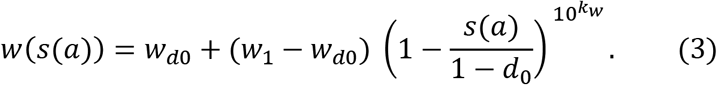

Here, *w*_*d*0_ and *w*_1_ are the minimum and maximum predator attack rates, while the parameter *k_w_* allows for flexibility in the predator attack rate with respect to the prey vividness; *k_w_* = 0 corresponds to a linear function, whereas *k_w_* < 0 and *k_w_* > 0 correspond to concave and convex forms of *w*(*s*(*a*)), respectively (Fig. 1c). Individual predators may differ in the parameter *k_w_*, thus representing diet breadth. The parameter *k_w_* thus determines predator phenotype, and we are interested in its evolution, too. The reason for considering 10^*k_w_*^ instead of just *k_w_* is the same as for the prey trait *k_d_*.

With this, the probability that a prey individual *j* escapes predation within a time step is

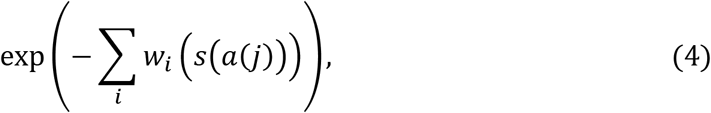

where the index *i* runs over all predators. The probability that when a prey individual *j* is consumed it is a predator *k* that consumes it, is

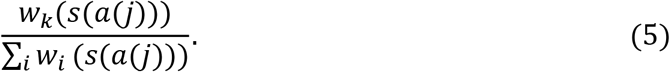

Once all prey individuals are tested for being consumed or not, the mean number of offspring a predator produces is calculated as proportional to the total energy obtained from prey consumption:

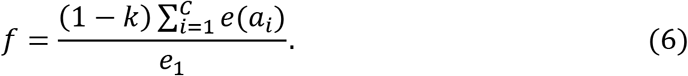

Here *C* is the number of prey individuals a predator consumes, *e*(*a_i_*) is an energy assimilated from the *i*-th consumed prey of age *a_i_*, *e*_1_ is energy needed to produce one predator offspring, and *k* is a proportion of the assimilated energy from food that goes to predator maintenance rather than reproduction. We assume for simplicity that *e*(*a*) = *e*_0_ + *k_e_a*, for some positive constants *e*_0_ and *k_e_*. Although it is likely that the prey energy *e*(*a*) eventually saturates with age, predation does not allow reaching high ages of prey and therefore our linear approximation appears to be a good approximation. The actual number of offspring born to a predator individual is assumed Poisson-distributed, with mean *f*. Individual predators suffer from background mortality such that each predator dies with probability *m* per time step.

Finally, we assume that the predator trait *k_w_* shaping the relationship between prey vividness and predator willingness to attack prey is traded off with another predator trait, so as to prevent runaway evolution to ever larger negative values of *k_w_* corresponding to the maximum predator attack rate on prey of any age and any mortality profile. Increasingly lower *k_w_* means that predators search for and attack increasingly more vivid prey (Fig. 1c) and need thus invest more to maintenance as opposed to reproduction. We model this as an increase in the maintenance parameter *k* with decreasing *k_w_*, and let

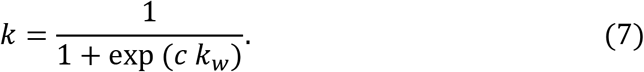

for some positive trade-off strength *c*.

### Predator-prey eco-evolutionary dynamics

Within each time step, prey first mate and reproduce. We assume that mates are chosen randomly, regardless of their phenotypes. Upon mating, a Poisson-distributed number of offspring are produced, with mean *b*(*x*, *k_d_*), where *k_d_* is the trait of a “mother” randomly chosen from the mating pair. The offspring are born with age 1 (as we emphasize earlier, fecundity already accounts for the first-year mortality). The phenotype *k_d_* of each prey offspring is then determined as follows. First, each offspring inherits the trait value from one of its parents, with equal probability. Then, with probability *p_m_* mutation occurs on this inherited trait. Upon mutation, a value generated from a normal distribution with zero mean and variance 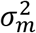 is added to the offspring’s trait value.

Background mortality of other prey than the offspring then occurs: individual prey die each with their respective probability *d*(*a*). This is followed by the extra mortality due to predation, described above and excluding just produced prey offspring. The age of all surviving prey individuals is then augmented by 1 and we record the trait distribution of the prey population (and calculate its mean and variance). Eventually, a maximum of *N* prey are randomly selected to form the population at the beginning of the next time step.

Each predator then mates and reproduces. Also here, mates are chosen randomly, regardless of their phenotypes. Upon mating, a Poisson-distributed number of offspring are produced, with mean *f*. If evolution works also on predators, the reproduction and phenotype *k_w_* of each predator offspring are determined analogously as in prey. Natural mortality of other predators than the offspring then follows. Finally, a maximum of *P* predators are randomly selected to form the population at the beginning of the next time step.

We note that we also considered a more complex model of predator and prey genetics, assuming that both *k_d_* and *k_w_* were polygenic traits, represented by a large number of haploid loci with additive allelic effects among loci (a variant of the procedure due to Holt, Gomulkiewicz, and Barfield^27^). The results of the model involving this quantitative genetic step were analogous to those produced by the simpler evolutionary model described above.

### Pairwise invasibility plots

Pairwise invasibility plot (commonly abbreviated to PIP) is a standard way of visualizing evolution of a single trait, assuming that the timescale at which ecological dynamics operate is much faster than the timescale of evolution.^28^ In its original form, it plots, for each pair of resident and mutant traits, fitness of a rare mutant entering a resident population at a stable ecological attractor (most commonly at a stable equilibrium). The idea is that the mutant either replaces the resident (if it has positive fitness) or fades away (when it has negative fitness). Evolution is then viewed as a sequence of mutations followed by replacements and PIPs are a way to visualize and follow such mutation-replacement dynamics.^28^ In particular, when mutations are small, evolution proceeds along the PIP diagonal, as any point on this diagonal stands for a resident population only. Invasion of mutants with a higher trait means moving the point vertically above the diagonal, and if such invasion is successful (trait replacement occurs) mutants become the new residents and we return to the PIP diagonal horizontally. If the invasion is not successful, we return to the PIP diagonal in the vertical direction, as nothing actually happened in the evolutionary sense. Analogously, invasion of mutants with a lower trait means moving the point vertically below the diagonal. In any case, another mutation is then assumed. This sequence of mutations and replacements stops either at an intermediate trait value or at a border of the adopted trait values interval.

The PIPs we present below are a stochastic variant of PIPs commonly presented in the literature, and are composed in the following way. For each selected combination of resident and mutant traits, we first let the resident population settle at its ecological attractor via running its dynamics for 1000 time steps. Then, we add a small number of mutants and follow resident-mutant competition dynamics for other 1000 time steps. We then record the proportion of mutants in the final population. Very low proportions thus mean mutants fade away, whereas proportions close to one indicate that mutants eventually replace residents. The proportion of mutants in the final population thus in our case plays the role of fitness in the original PIP construction.

## Results

### Evolution of prey aging rate in the absence of predators

We start with exploring the evolution of prey aging rate in the absence of predators. This represents the baseline scenario with which we compare our other results. When there is no cost of aging (*x* = 0), prey age at an increasingly lower rate, as *k_d_* steadily increases, but there is no apparent lower limit on the aging rate (Fig. 2, top row).

**Figure 2:**
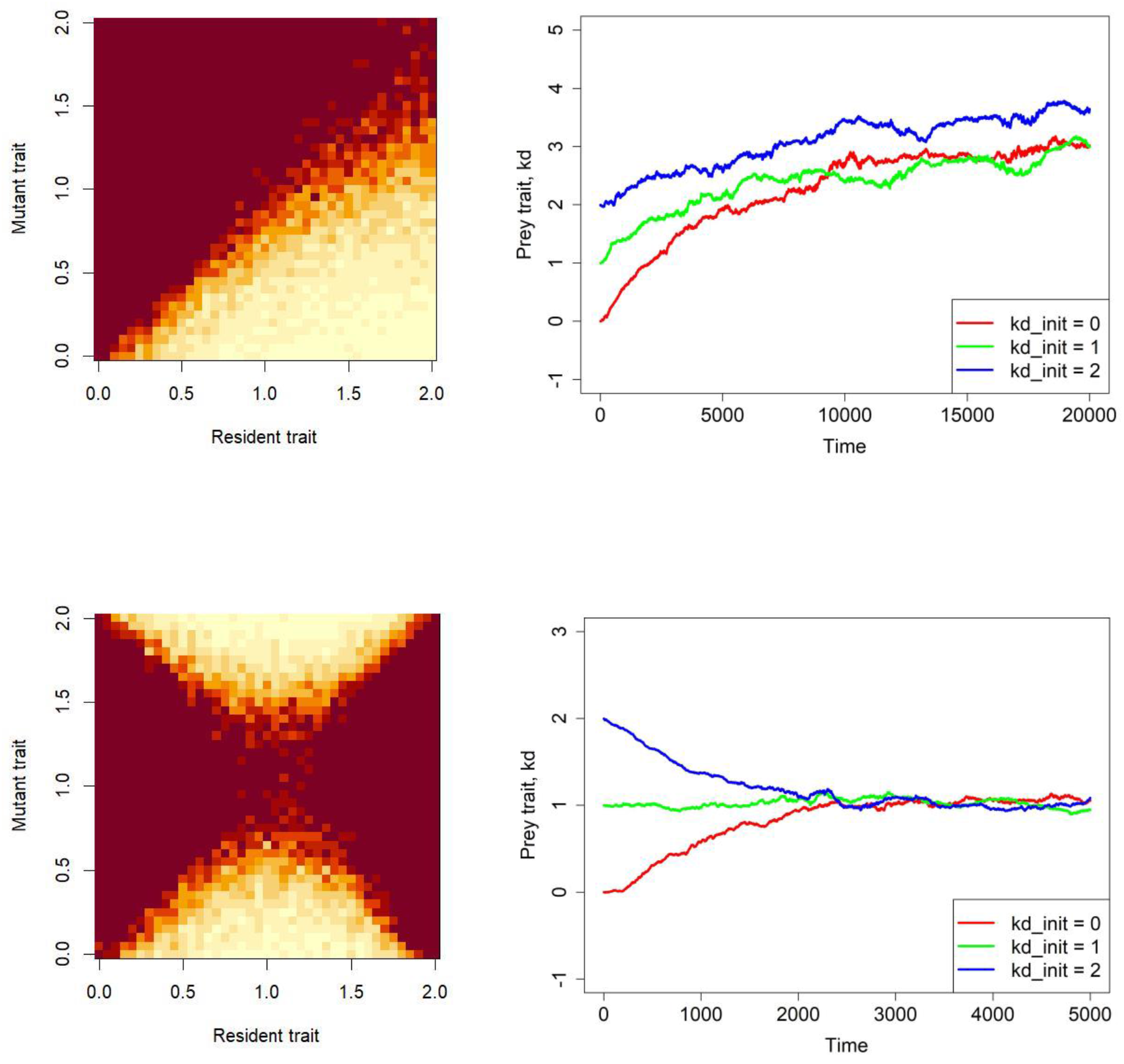
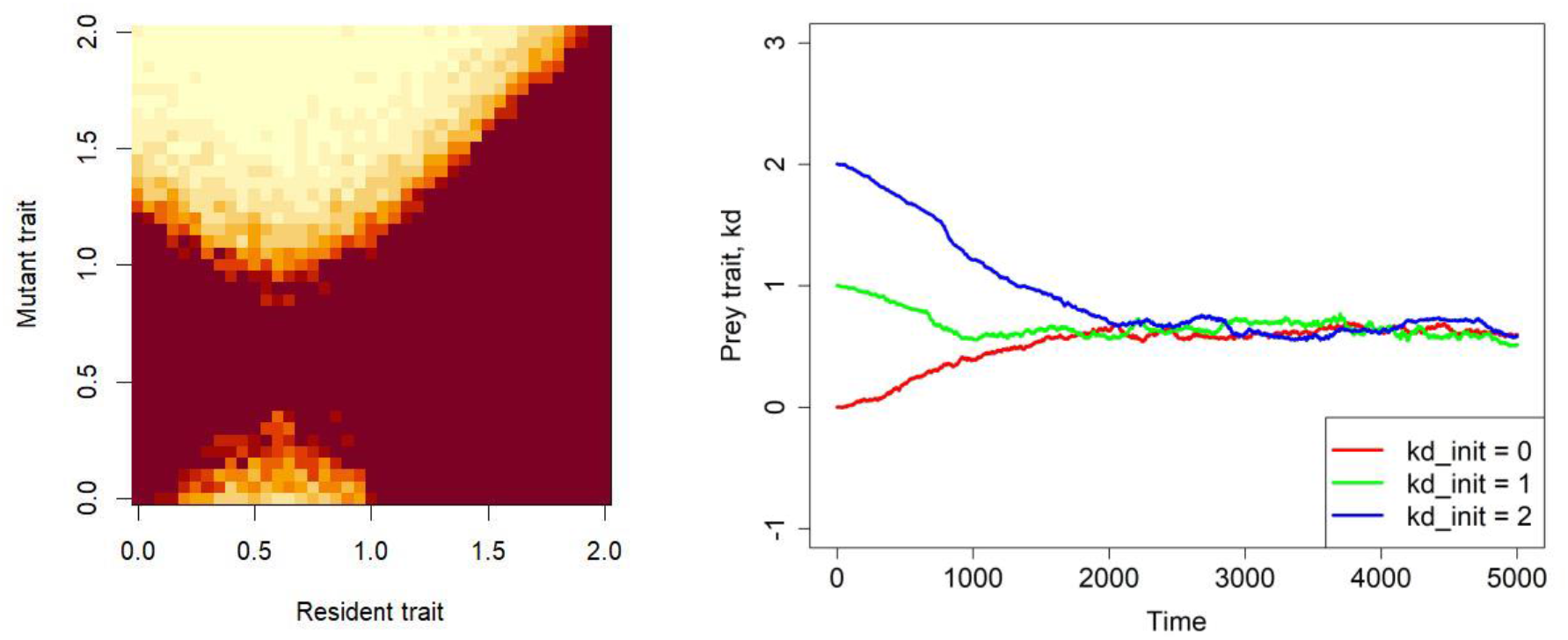
Evolution of the prey trait *k_d_* in the absence of predators when there is no cost of aging (*x* = 0, top row), a low cost (*x* = 0.01, middle row) or a high cost (*x* = 0.05, bottom row). The left panels are the respective pairwise invasibility plots: whereas bright colors correspond to low proportions of mutants in the population and hence indicate mutant extinction, dark colors correspond to high proportions of mutants in the population and hence indicate that mutants eventually replace residents. The right panels show one replicate of the temporal course of evolution for three different initial mean values of *k_d_*; mean *k_d_* over the prey population is shown. Other parameters are as in Table 1.

When a cost of aging is present, the prey trait *k_d_* appears to stabilize (Fig. 2, middle row). Hence, there appears to be an optimal aging rate modulated by the aging rate-fecundity trade-off. Indeed, *k_d_* attains lower values by evolution (that is, faster aging occurs) when the aging cost is higher (Fig. 2, bottom row) or when the prey birth rate is higher (Fig. 3, showing also the case when both the aging cost and the prey birth rate are simultaneously higher).

**Figure 3:**
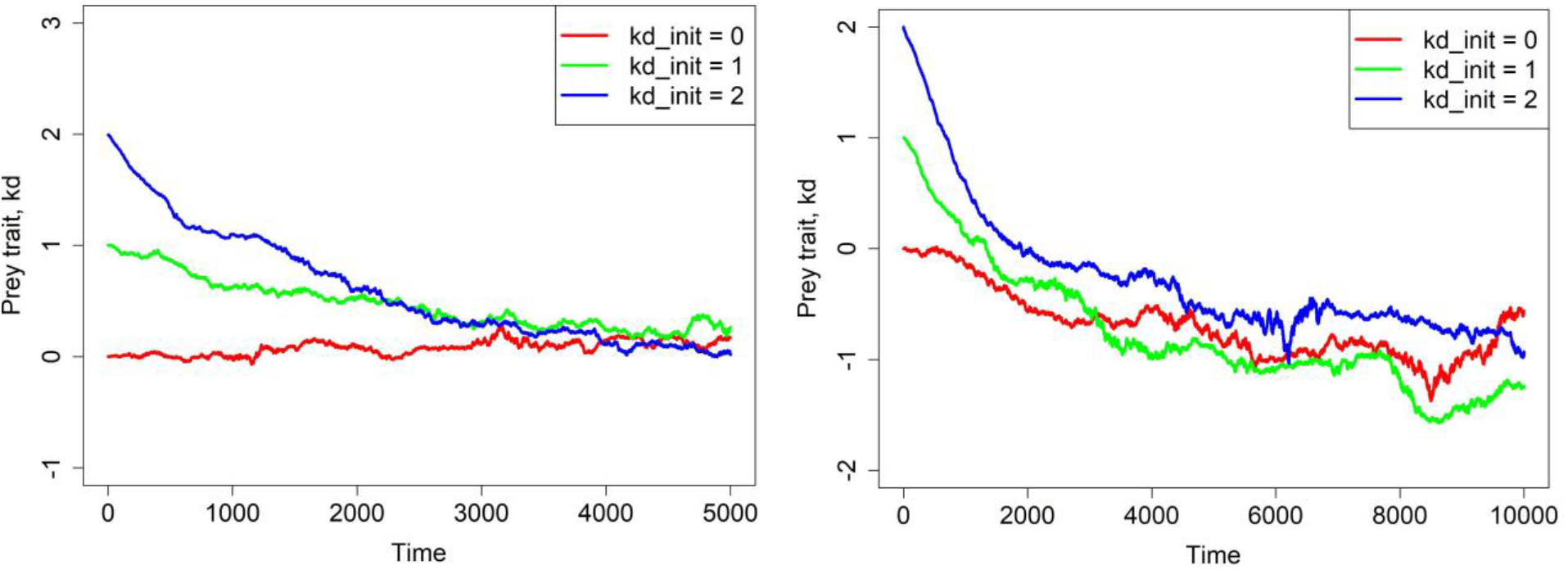
Evolution of the prey trait *k_d_* in the absence of predators when aging bears a cost. Left panel: *x* = 0.01 and the value of *b*_0_ increased from 1 to 5. Right panel: the aging cost increased to *x* = 0.05 and the value of *b*_0_ increased from 1 to 5. Other parameters are as in Table 1. Both panels show one replicate of temporal course of evolution for three different starting mean values of *k_d_*; mean *k_d_* over the prey population is shown.

### Evolution of prey aging rate in the presence of predators

We now examine the evolution of prey aging rate in the presence of predators. The aging rates attained by evolution in the previous section are the optimal aging rates set by particular strengths of the aging rate-fecundity trade-off, and therefore we expect that with predators prey cannot evolve slower aging rate than without predators. Rather, we expect that predator presence will lead to faster aging, depending on predator characteristics.

Figure 4 shows an effect of a different number of predators and of a different predator trait value *k_w_*. We note that predators with lower *k_w_* include into their diet more vivid (thus relatively younger) prey individuals. If *k_w_* is relatively large so that only relatively older individuals form predator diet, the optimal prey strategy does not differ too much from that without predators, which is to age relatively slowly close to the lower bound set by the aging rate-fecundity trade-off. On the other hand, when predators consume also relatively young prey (such as when they have negative *k_w_*), the optimal prey strategy appears to be to age faster and rather produce as many offspring as possible as soon as they can (so lower *k_d_* evolves; Fig. 4). Moreover, while the number of predators does not appear to affect results of evolution when *k_w_* is relatively large, lower values of *k_d_* are attained if there are more predators around when *k_w_* is relatively small (Fig. 4).

**Figure 4:**
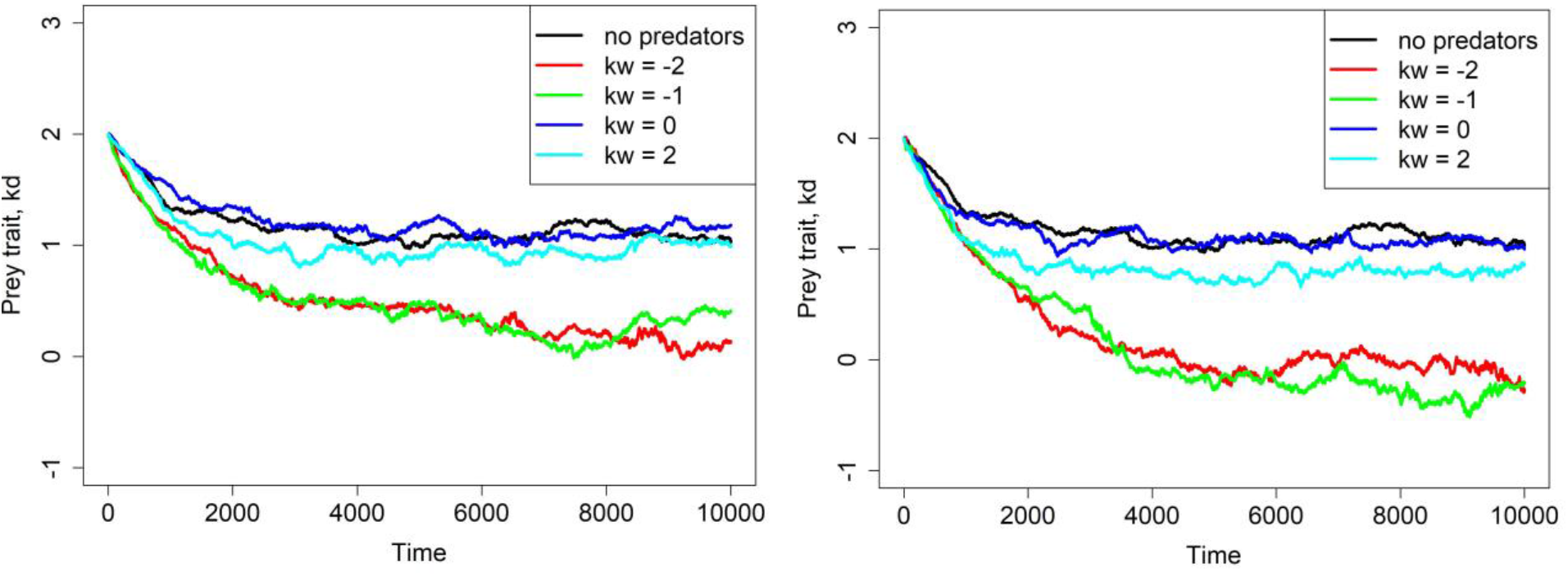
Evolution of the prey trait *k_d_* in the presence of predators under several fixed values of predator trait *k_w_*. The left panel is for 200 predators, the right one is for 500 predators. One replicate of temporal course of evolution for each value of *k_w_* is shown; the initial mean value of *k_d_* in prey is always set to 2. Other parameters are as in Table 1. Predators were not allowed to evolve and all has the same value of *k_w_*. The no-predator scenario is shown in black.

### Coevolution

Finally, we study coevolution between the prey trait *k_d_* and the predator trait *k_w_*. Interestingly, the results differ substantially for different starting values of the predator trait *k_w_* (Fig. 5). Consider first the left panel of this figure. Negative staring values of *k_w_* lead to evolution of low values of *k_d_*, since prey need to produce as many offspring as possible as fast as possible and slower aging thus has no obvious advantage (see Fig. 4). We recall that the lower is *k_w_* the more predators add to their diet more vivid (thus relatively younger) prey (see Fig. 1c). When *k_d_* becomes low and prey thus age faster, so becomes low also vividness of most prey individuals. As a consequence, it is no more advantageous for predators to have low *k_w_* which means high maintenance *k*. Selection thus prefers higher values of *k_w_* which therefore starts to rise. When *k_w_* increases, it becomes advantageous for prey to increase *k_d_* and so its vividness at any age; *k_d_* thus increases, eventually reaching a value close to the upper bound set by the aging rate-fecundity trade-off in prey. Now consider the right panel of Fig. 5. Relatively large starting values of *k_w_* make the prey trait *k_d_* quickly attain a value bounded by the aging rate-fecundity trade-off in prey and *k_w_* then attains an optimal value for that *k_d_*.

**Figure 5:**
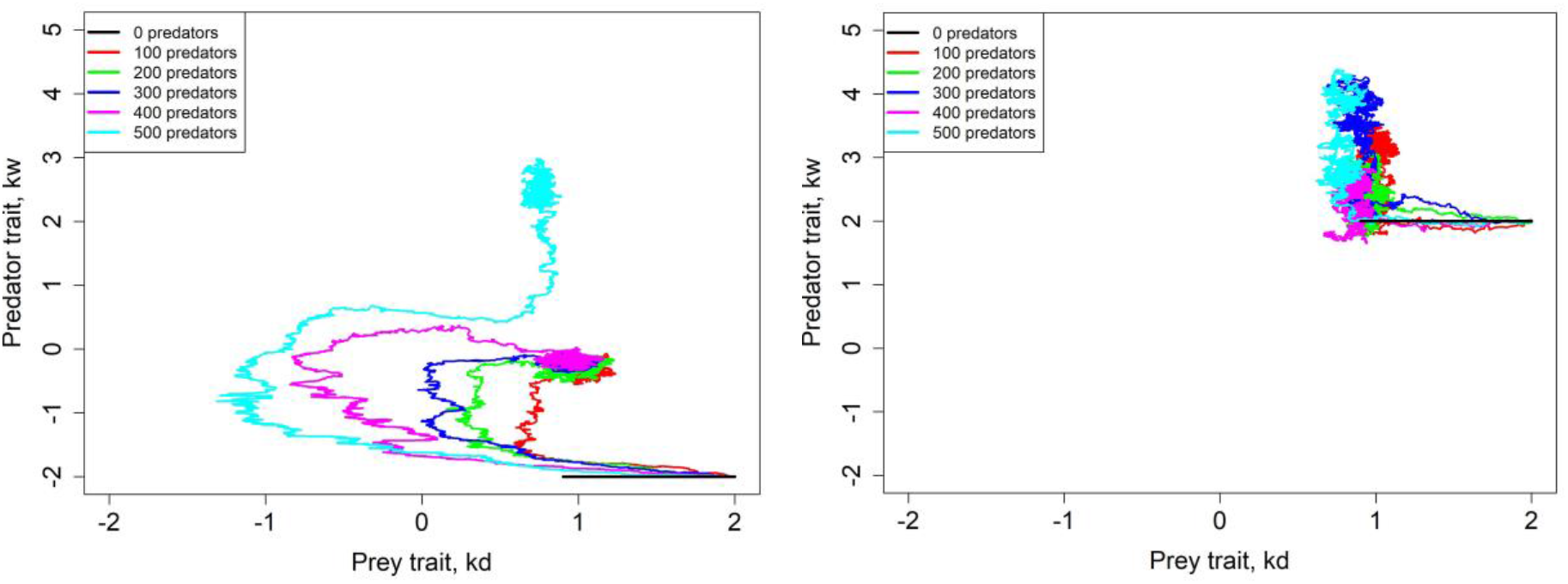
Coevolution of the prey trait *k_d_* and the predator trait *k_w_*. One replicate of temporal course of coevolution is shown for diverse numbers of predators, starting with two different values of *k_w_*. Other parameters are as in Table 1. The no-predator scenario is shown in black (horizontal black line going from *k_d_* = 2 to *k_d_* slightly lower than 1). The simulation is run for 40000 time steps.

### No aging rate-fecundity trade-off

Recently, some researchers have argued against aging rate-fecundity trade-offs as drivers of senescence and life span evolution.^29^ To respect these views, we have also conducted simulation experiments with predators but without the aging rate-fecundity trade-off in prey (i.e. *x* = 0).

Interpretation of results, in this case, is not as clear as under the aging rate-fecundity trade-off in prey (Fig. 6). For example, some trajectories in Fig. 6 suggest that the presence of predators may speed up the evolution of slower aging rate in prey (*k_w_* = −1, left panel), but some contrarily suggest stabilizing selection (*k_w_* = −2, right panel). Such an ambiguity also remains with regards to the coevolution of prey and predator traits *k_d_* and *k_w_* (Fig. 7).

**Figure 6:**
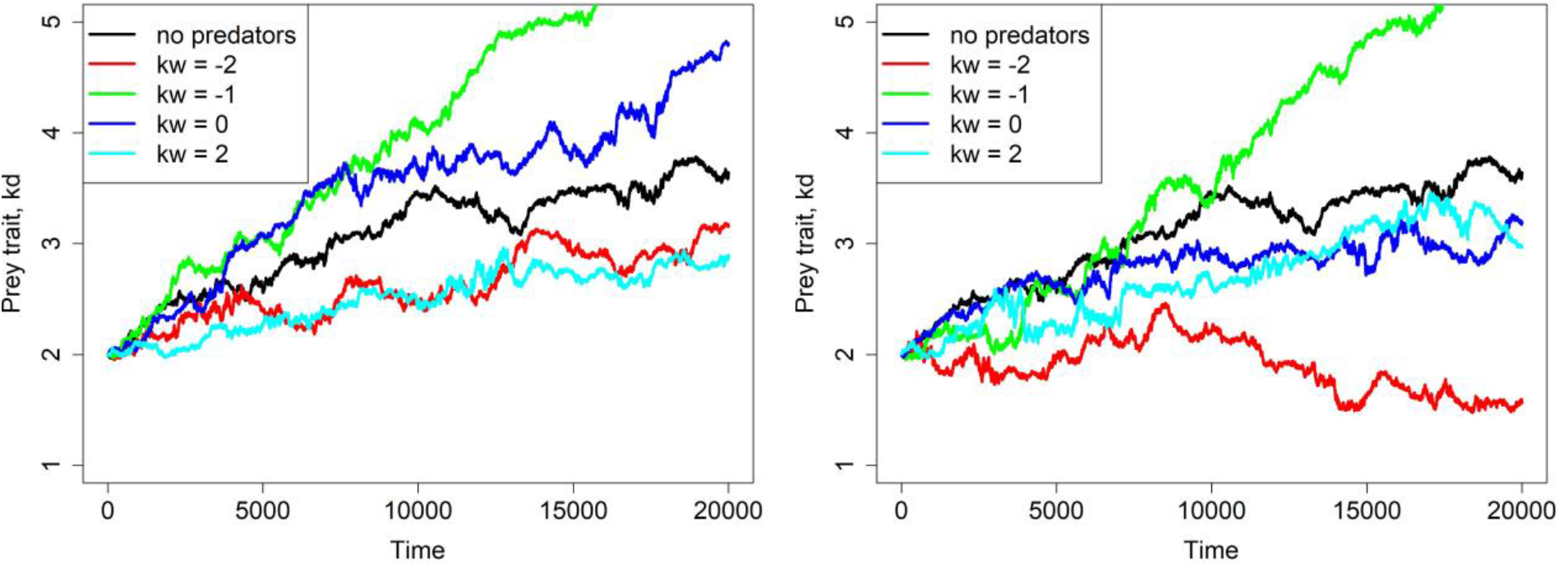
Evolution of the prey trait *k_d_* in the absence of aging rate-fecundity trade-off (*x* = 0) yet in the presence of predators, under several fixed predator traits *k_w_*. The left panel is for 200 predators, the right one is for 500 predators. One replicate of temporal course of evolution for each value of *k_w_* is shown. Other parameters are as in Table 1. Predators were not allowed to evolve and all has the same value of *k_w_*. The no-predator scenario is shown in black.

**Figure 7:**
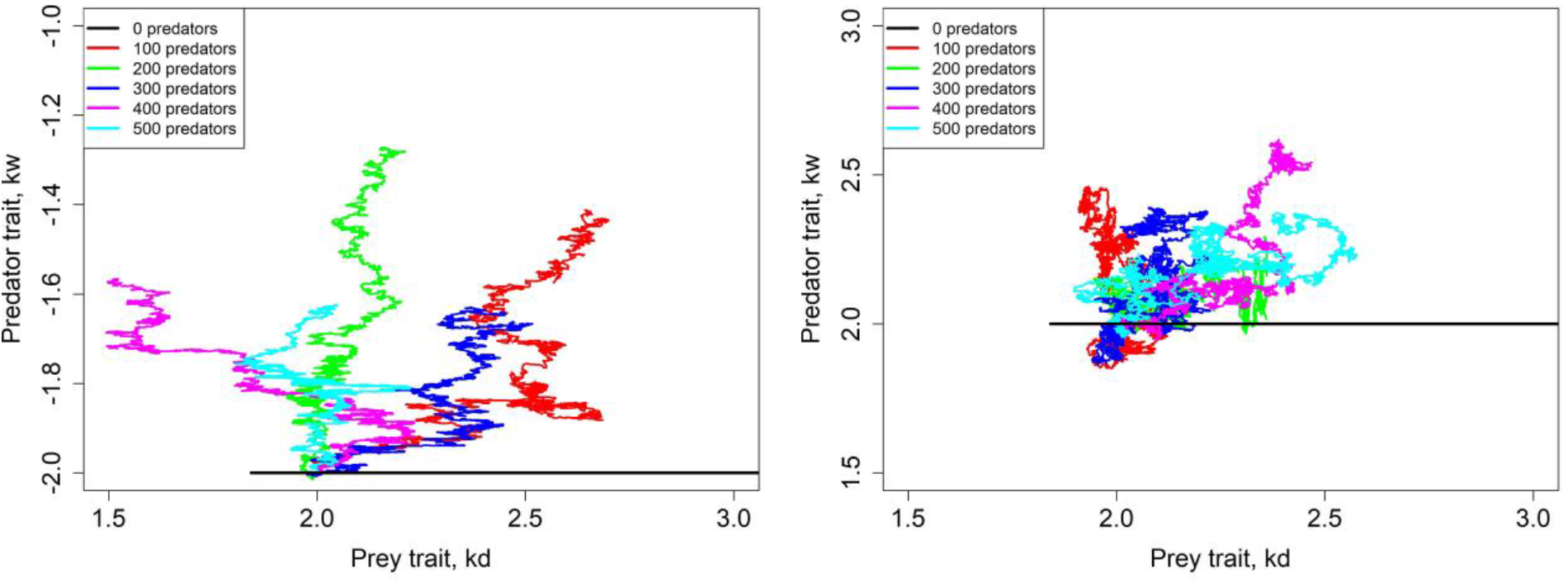
Coevolution of the prey trait *k_d_* and the predator trait *k_w_* in the absence of aging rate-fecundity trade-off (*x* = 0). One replicate of temporal course of coevolution is shown for diverse numbers of predators, starting with two different values of *k_w_*. Other parameters are as in Table 1. The no-predator scenario is shown in black (horizontal black line starts at *k_d_* = 2 and eventually grows; it is actually the black curve from Fig. 6). The simulation is run for 40000 time steps.

## Discussion

To summarize, our results show that if a trade-off between the pace of aging and fecundity exists in a given species, then this trade-off is the main factor setting the course of evolution of aging. Predation then affects the evolution of the pace of aging only if there is no significant preference for relatively older prey. However, even then, the coevolution of predators and prey leads, in the long run, to the same pace of aging in prey as evolution without predators. On the other hand, our simulations also show that when the trade-off between aging and fecundity (in prey) is not present, the pace of aging evolves progressively slower without any lower limit. Furthermore, without the trade-off between aging and fecundity, predation becomes much more important and can lead to the evolution of both faster and slower aging, apparently at random. The most likely explanation of this effect seems to be that predation lowers the effective population size and thus, amplifies the effect of genetic drift.

Results of our simulations contradict a long-standing theoretical prediction that populations experiencing higher extrinsic mortality should always evolve to age faster.^8,30^ More importantly, they help to explain some previous experimental findings, e.g., that natural populations of *Daphnia ambigua* living in lakes varying dramatically in the intensity and duration of predation, age at the same pace.^31^ Or that natural populations of guppies experiencing higher mortality do not evolve earlier set of senescence with regards to mortality or reproduction.^32^ Furthermore, at least three testable predictions may be derived from the results of our simulations.

First, there is no trade-off between aging rate and fecundity in organisms with negligible senescence. In some organisms, this prediction may be challenging to test. However, in case of hydra, the absence of trade-off between aging rate and fecundity seems to be almost certain. Hydra can live for hundreds or even thousands of years^2,3^, while its mortality and fertility are constant^3,4^. If there was a trade-off between aging rate and fecundity in hydra it should reproduce particularly slowly if at all. This is, however, not the case. Thus, such trade-off in hydra most likely does not exist or is minuscule. Second, because the pace of aging varies considerably between species, the strength of trade-off between aging rate and fecundity must also vary considerably between different species and should be malleable by evolutionary forces as well. Third, interventions aiming at the trade-off between aging rate and fecundity should be especially potent ways how to slow-down aging. This prediction is in a good agreement with a long-known fact that ablation of germline can, at least in some organisms, prolong life-span^33,34^. However, our results suggest that the optimal approach to slow down aging should aim at modulating the pathways/signals regulating the trade-off between cellular maintenance and reproduction instead of focusing on one of its parts, i.e., germline.

Overall, the results presented here provide a novel insight into the evolution of aging. They suggest an easy to grasp explanation of the great diversity of aging found in nature, including its absence, without a need to invoke controversial concepts such as programed aging. Moreover, testable predictions based on these results translate into exciting opportunities for future research.

## Acknowledgments

The project was supported by the CETOCOEN PLUS (CZ.02.1.01/0.0/0.0/15_003/0000469) project of the Ministry of Education, Youth and Sports of the Czech Republic. The project was also supported by CETOCOEN EXCELLENCE Teaming 2 project supported by Horizon2020 (857560) and the Ministry of Education, Youth and Sports of the Czech Republic (02.1.01/0.0/0.0/18_046/0015975). As well as by the RECETOX Research Infrastructure (LM2018121). Furthermore, P.L. received support from the Brno Ph.D. Talent competition.

## Authors’ Contribution

P.L. formulated the research problem, interpreted the results, designed the simulation scenarios, and co-wrote the manuscript. J.B.V. supervised the project and obtained funding. L.B. developed and coded the mathematical model, interpreted the results, designed the simulation scenarios and co-wrote the manuscript. All authors reviewed the manuscript.

## Data availability

The datasets generated and/or analyzed during the current study are available from the corresponding author upon reasonable request.

## Code availability

The computer code used to generate results reported in this article is available from the corresponding author upon reasonable request, and will be released at publication of this article.

## Conflict of interest

The authors declare no conflict of interest.

## Notes

### Competing Interest Statement

The authors have declared no competing interest.

## References

1. Kirkwood, T. B. L. & Austad, S. N. Why do we age? Nature 408, 233 (2000).

2. Jones, O. R. et al. Diversity of ageing across the tree of life. Nature 505, 169–173 (2014).

3. Schaible, R. et al. Constant mortality and fertility over age in Hydra. Proc. Natl. Acad. Sci. 112, 15701–15706 (2015).

4. Martínez, D. E. Mortality patterns suggest lack of senescence in hydra. Exp. Gerontol. 33, 217–225 (1998).

5. Ruby, J. G., Smith, M. & Buffenstein, R. Naked mole-rat mortality rates defy Gompertzian laws by not increasing with age. eLife 7, e31157 (2018).

6. Baudisch, A. The pace and shape of ageing. Methods Ecol. Evol. 2, 375–382 (2011).

7. Kirkwood, T. B. L. & Melov, S. On the Programmed/Non-Programmed Nature of Ageing within the Life History. Curr. Biol. 21, R701–R707 (2011).

8. Williams, G. C. Pleiotropy, Natural Selection, and the Evolution of Senescence. Evolution 11, 398–411 (1957).

9. Kirkwood, T. B. L. Evolution of ageing. Nature 270, 301 (1977).

10. Kirkwood T. B. L., Holliday Robin, Maynard Smith J. & Holliday Robin. The evolution of ageing and longevity. Proc. R. Soc. Lond. B Biol. Sci. 205, 531–546 (1979).

11. Jones, O. R. & Vaupel, J. W. Senescence is not inevitable. Biogerontology 18, 965–971 (2017).

12. Longo, V. D., Mitteldorf, J. & Skulachev, V. P. Programmed and altruistic ageing. Nat. Rev. Genet. 6, 866–872 (2005).

13. Mitteldorf, J. & Martins, A. C. R. Programmed life span in the context of evolvability. Am. Nat. 184, 289–302 (2014).

14. Libertini, G. An adaptive theory of increasing mortality with increasing chronological age in populations in the wild. J. Theor. Biol. 132, 145–162 (1988).

15. Bowles, J. T. The evolution of aging: a new approach to an old problem of biology. Med. Hypotheses 51, 179–221 (1998).

16. Bredesen, D. E. The non-existent aging program: how does it work? Aging Cell 3, 255–259 (2004).

17. Prinzinger, R. Programmed ageing: the theory of maximal metabolic scope. EMBO Rep. 6, S14–S19 (2005).

18. Mitteldorf, J. & Goodnight, C. Post-reproductive life span and demographic stability. Oikos 121, 1370–1378 (2012).

19. Kowald, A. & Kirkwood, T. B. L. Can aging be programmed? A critical literature review. Aging Cell 15, 986–998 (2016).

20. Travis, J. M. J. The Evolution of Programmed Death in a Spatially Structured Population. J. Gerontol. Ser. A 59, B301–B305 (2004).

21. Mitteldorf, J. & Pepper, J. Senescence as an adaptation to limit the spread of disease. J. Theor. Biol. 260, 186–195 (2009).

22. Martins, A. C. R. Change and Aging Senescence as an Adaptation. PLOS ONE 6, e24328 (2011).

23. Werfel, J., Ingber, D. E. & Bar-Yam, Y. Programed Death is Favored by Natural Selection in Spatial Systems. Phys. Rev. Lett. 114, 238103 (2015).

24. Lenart, P. & Bienertová-Vašků, J. Keeping up with the Red Queen: the pace of aging as an adaptation. Biogerontology 18, 693–709 (2017).

25. Lenart, P., Bienertová-Vašků, J. & Berec, L. Evolution favours aging in populations with assortative mating and in sexually dimorphic populations. Sci. Rep. 8, 16072 (2018).

26. Harb, S. C. et al. Estimated age based on exercise stress testing performance outperforms chronological age in predicting mortality. Eur. J. Prev. Cardiol. 2047487319826400 (2019) doi:10.1177/2047487319826400.

27. Holt, R. D., Gomulkiewicz, R. & Barfield, M. The phenomenology of niche evolution via quantitative traits in a ‘black-hole’ sink. Proc. Biol. Sci. 270, 215–224 (2003).

28. Brännström, Å., Johansson, J. & Von Festenberg, N. The Hitchhiker’s Guide to Adaptive Dynamics. Games 4, 304–328 (2013).

29. Cohen, A. A., Coste, C. F. D., Li, X.-Y., Bourg, S. & Pavard, S. Are trade-offs really the key drivers of ageing and life span? Funct. Ecol. 34, 153–166 (2020).

30. Medawar, P. B. An unsolved problem of biology. (Published for the College by H.K. Lewis, 1952).

31. Walsh, M. R., Whittington, D. & Walsh, M. J. Does variation in the intensity and duration of predation drive evolutionary changes in senescence? J. Anim. Ecol. 83, 1279–1288 (2014).

32. Reznick, D. N., Bryant, M. J., Roff, D., Ghalambor, C. K. & Ghalambor, D. E. Effect of extrinsic mortality on the evolution of senescence in guppies. Nature 431, 1095–1099 (2004).

33. Hsin, H. & Kenyon, C. Signals from the reproductive system regulate the lifespan of C. elegans. Nature 399, 362–366 (1999).

34. Antebi, A. Regulation of longevity by the reproductive system. Exp. Gerontol. 48, 596–602 (2013).

